# ProtRNA: A Protein-derived RNA Language Model by Cross-Modality Transfer Learning

**DOI:** 10.1101/2024.09.10.612218

**Authors:** Ruoxi Zhang, Ben Ma, Gang Xu, Jianpeng Ma

## Abstract

Protein language models (PLM), such as the highly successful ESM-2, have proven particularly effective. However, language models designed for RNA continue to face challenges. A key question is: can the information derived from PLMs be harnessed and transferred to RNA? To investigate this, a model termed ProtRNA has been developed by cross-modality transfer learning strategy for addressing the challenges posed by RNA’s limited and less conserved sequences. By leveraging the evolutionary and physicochemical information encoded in protein sequences, the ESM-2 model is adapted to processing “low-resource” RNA sequence data. The results show comparable or superior performance in various RNA downstream tasks, with only 1/8 the trainable parameters and 1/6 the training data employed by the primary reference baseline RNA language model. This approach highlights the potential of cross-modality transfer learning in biological language models.

## Introduction

As the Transformer architecture continues to dominate state-of-the-art (SOTA) performance in natural language processing (NLP) tasks, its applications to biological sequences have gained increasing attention. By treating the building blocks of biological sequences as “tokens”, researchers have developed large Biological Language Models (BLM) pretrained on DNA, RNA, or protein sequence data. These models are being deployed for a wide range of structural and functional prediction tasks.

BERT-style [1] BLMs, pretrained by Masked Language Modeling (MLM) and can produce bi-directional representations, have recently gained unignorable advancements. Notably, BERT-style protein language models, like ESM-2, have excelled in structure prediction [2] and various other downstream tasks. These models are thought to internalize evolutionary patterns by pretraining on large protein sequence databases [3]. The bidirectional self-attention mechanism in these models is suggested to enable the capture of residue-residue contacts, storing this information in attention heads and representations [4].

Unlike protein language models, such as SOTA ESM-2, few general-purpose RNA language models show robust performance. RNA-FM [5] is one of the first RNA foundation language models trained on noncoding RNA (ncRNAs). It consists of 12 standard encoder transformer blocks, with 640 embedding dimensions, and is trained on 23 million sequences from the RNAcentral database [6]. RiNALMo [7] is a larger model with 33 transformer blocks and 1280 embedding dimensions and is trained on a curated 36 million ncRNA dataset, with some architectural modifications incorporating rotary positional embedding (RoPE) and SwiGLU activation function. RNAErnie [8] introduces a multilevel masking strategy for pretraining, including masking at the base, subsequence, and motif levels. Like RNA-FM, RNAErnie is trained on ncRNA sequences from RNAcentral and features a medium-sized architecture with 12 transformer blocks and 768 embedding dimensions.

Recent notable advances in RNA language models utilized RNA sequences other than ncRNAs and architectures extending beyond BERT. GARNET [9] is a comprehensive database containing over 400,000 bacterial and archaeal genomes from the Genome Taxonomy Database (GTDB). A GPT-style autoregressive model was proposed to be trained on the database with triplet nucleotide tokenization and structural-aware graph neural networks (GNNs). LARMAR [10] uses 15 million RNA sequences of mammalian and viral genes and transcripts to build a foundational model for RNA regulation. Orthrus [11] adopts a biological augmented contrastive learning objective in a Mamba-based state-space model, trained on curated pairs of RNA transcripts, generating functional mRNA embeddings relevant to properties like subcellular localization.

Nonetheless, the effectiveness of language modeling, instead of the Multiple Sequence Alignment (MSA) approach, in capturing evolutional information in RNA has been questioned [12]. RNA language modeling is considered intrinsically more challenging than protein due to the limited four-letter alphabet and the lower conservation of RNA sequences [13]. This results in sparse and noisy data, making RNA sequences more difficult to model effectively. The low quality of data in RNA language thus renders it a “low-resource” language in NLP terminology, in contrast to the ‘resource-rich’ protein language.

Parameter-efficient fine-tuning (PEFT) [14] enables large pre-trained models to be adapted to shifted distributions by updating a small fraction of parameters. Methods such as Low-Rank Adaptation (LoRA) [15] and its generalized variants, such as GLoRA [16] and PRoLoRA [17], accomplish this through low-rank matrix or dynamic adapters, usually modifying less than 1% of the total model parameters. Another common strategy for efficient transfer learning is the partial unfreezing of transformer layers critical to the target distribution [18] [19], which can be valuable for tasks that diverge from the original training distribution or involve new input modalities. Following the line of cross-lingual transfer learning studies in the field of NLP [20], we explored a cross-modality transfer learning pretraining strategy that leverages the SOTA ESM-2 model trained on the ‘resource-rich’ language protein sequences and adapts it to ‘low-resource’ ncRNA data (**Figure 1.a**).

**Figure 1.**
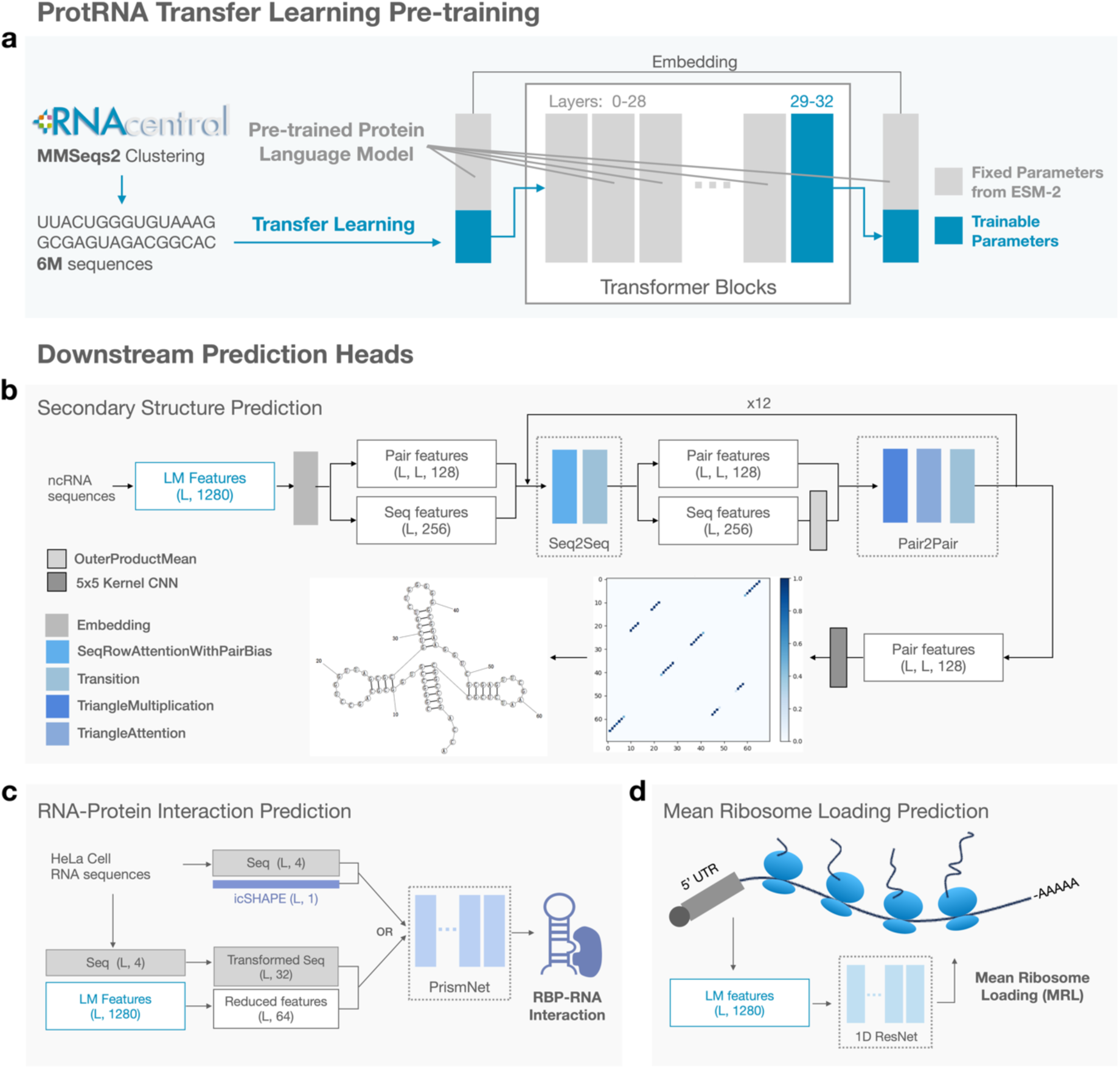
Overview of the ProtRNA transfer-learning pretraining strategy for RNA language model and downstream task architectures. a, the ProtRNA model utilizes initial weights from the SOTA protein language model ESM-2. During the cross-modality transfer learning phase, the model is pretrained on 6 million non-coding RNA sequences using the MLM task. It freezes the first 29 transformer layers, updating only RNA vocabulary embeddings and the last four layers (30-33). b-d, after pretrained for six epochs on RNA data, the ProtRNA model is evaluated on three downstream RNA tasks. For these tasks, task-specific heads receive sequence representations from the pretrained language model, and only the heads are further trained on respective training sets.

Our approach leverages the evolutionary priors and shared biophysical principles encoded in the pretrained protein ESM-2 model to facilitate RNA language modeling. In particular, this cross-modality transfer learning strategy–adapting a protein language model to RNA tasks–gains from partially unfreezing key transformer layers to capture RNA-specific structural and functional motifs. The ProtRNA model, derived from this transfer learning pretraining strategy of partial unfreezing deep layers, exhibits performance comparable with or superior to existing foundational RNA language models across various RNA downstream tasks (**Figure 1.b-d**). Remarkably, this was achieved with merely 1/8 trainable parameters and 1/6 training data employed by the primary reference model RiNALMo, underscoring the efficiency and effectiveness of our approach.

## Results

### Cross-Modality Transfer Learning Strategy for ProtRNA Pretraining

#### Parameter and Data Efficient Pretraining

Utilizing the ESM-2 model with 1280 embedding dimensions and 33 transformer layers, we froze the first 29 layers and trained only the final four layers and the embeddings for RNA tokens. The training was performed on 6 million representative RNA sequences clustered by MMSeqs2 [21] from the RNAcentral database. As seen from **Table 1**, compared with other BERT-style RNA language models, both the trainable parameters and the effective training data utilized in our pretraining process are considerably fewer. We will show in the following section that even with this reduction in computational resources, the performance of our model in various downstream tasks is comparable with or even superior to other models. The efficiency of our transfer learning method is demonstrated by the ablation study on a smaller dataset, shown in **Figure S1.a**, where the training loss and validation accuracy of the transfer learning method improves significantly faster than training from scratch with the same number of trainable parameters and randomly initialized weights. These findings are further supported by experiments using models trained on the full pretraining data (**Figure S1.b**) and the performance gap on the key downstream task of secondary structure prediction (**Table S1**). Overall, pretraining on ESM-2 weights consistently outperforms both the randomly initialized model and the model fine-tuned via LoRA.

**Table 1.** Comparison of Training Process and Model Architecture between ProtRNA and other foundational RNA language models. The dash (‘-’) indicates the information is not disclosed in the publication. The Uni-RNA model, not released publicly, is excluded from subsequent downstream task testing.

|  | ProtRNA | RiNALMo | Uni-RNA | RNA-FM | RNAErnie |
| --- | --- | --- | --- | --- | --- |
| Trainable Params. | 80M | 650M | 400M | 100M | 105M |
| Training Data | 6M | 36M | 500M | 23M | 23M |
| Total Params. | 650M | 650M | 400M | 100M | 105M |
| Total Layers | 33 | 33 | 24 | 12 | 12 |
| Embedding Dim. | 1280 | 1280 | 1280 | 640 | 768 |

#### Distinguishable Clustering of RNA Families and Protein-RNA Segregation in Representation Space

Figure 2 presents a t-SNE-based 2D visualization of the RNA and protein sequence representations generated by our final ProtRNA model. The RNA sequences are from the ArchiveII dataset [22], and the protein sequences are from UniProt [23]. The t-SNE algorithm effectively captures the local and global structure of the high-dimensional sequence data in a lower-dimensional space [24]. The visualization reveals that RNA sequences from different families form distinct clusters. The protein representations are also clearly segregated from the RNA representations, residing in the upper left corner of the plot. This outcome highlights the model’s ability to discriminate between RNA families while effectively distinguishing protein from RNA representations. Some families of ncRNA sequences are clustered in a similar region on the middle left of the t-SNE plot, particularly those with small sample sizes, such as telomerase, rRNA 16s, and grp2. This clustering may be attributed to the limited number of sequences in the test set and their underrepresentation in the pretraining data, which may hinder the model’s ability to distinguish these RNA families effectively.

**Figure 2.**
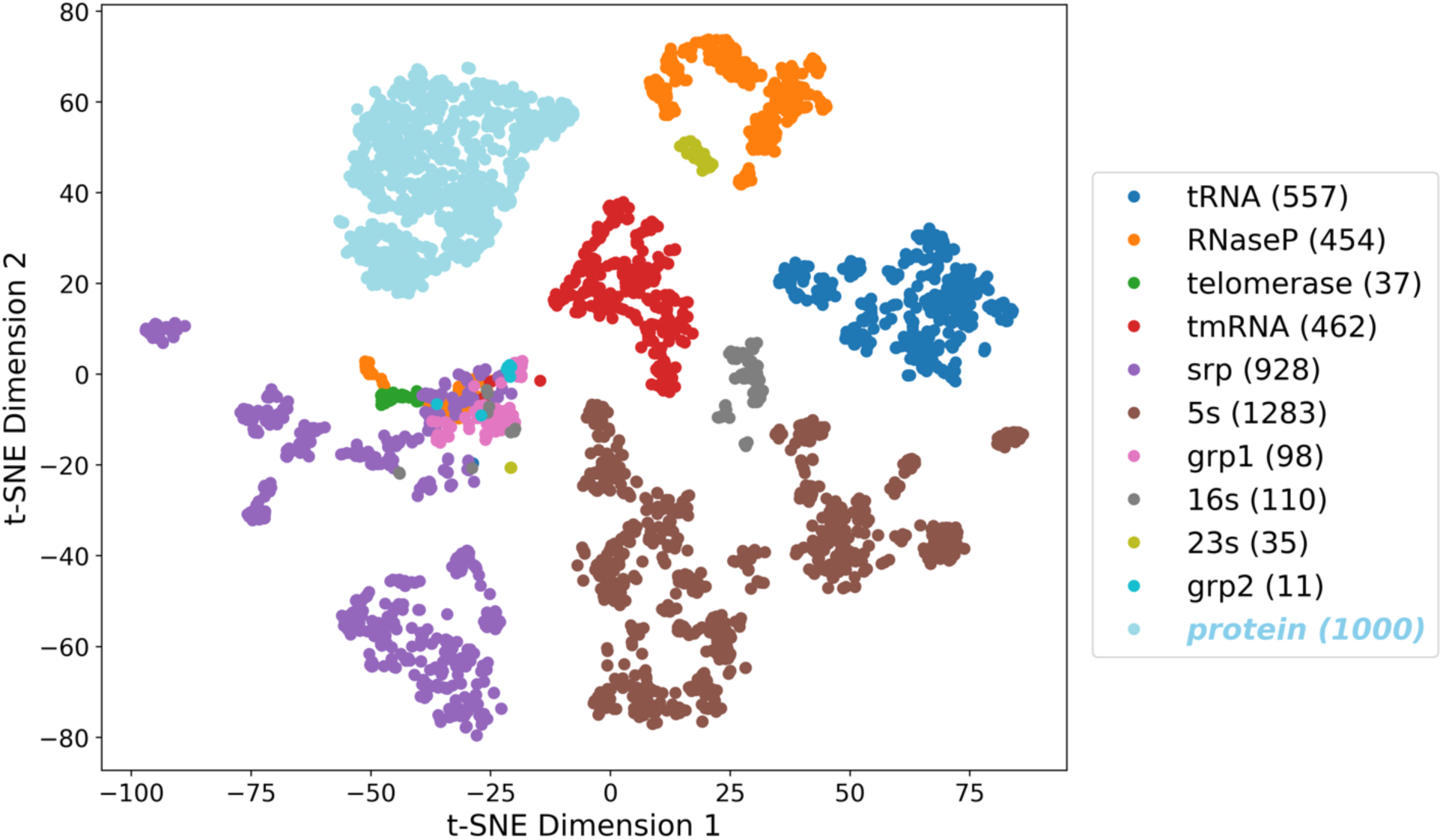
t-SNE visualization of ProtRNA Representations. RNAs from different families are colored differently. The protein cluster (denoted in sky blue) is distinctly separated from the RNA families at the upper left corner. The legend shows the names of the ncRNA families, with the number of sequences for each family from the ArchiveII dataset indicated in parentheses. For protein, the number of randomly selected sequences from UniProt is shown.

#### Distinctive Attention Pattern in the ProtRNA Model Where a Specialized Structure Head Emerges

Though the interpretability of BERT models in NLP, particularly how they encode dependency relations through self-attention, has been extensively studied [25] [26] [27] [28], research on the inner workings of BERT models in the context of biological sequences is relatively sparse. The multi-head self-attention mechanism is considered crucial, as it enables the model to capture pairwise dependencies within a sequence, especially long-range interactions, which may correspond to the spatial contacts that occur when a sequence folds into a stable three-dimensional structure [29]. Hereby we provide an attention analysis of our ProtRNA model and two other baseline RNA BERT models, with regard to two structural properties of RNA, applying the method devised by [4]. The visualization of the attention analysis results is presented in Figure 3. See **Methods** for details of the metric and the datasets used.

**Figure 3.**
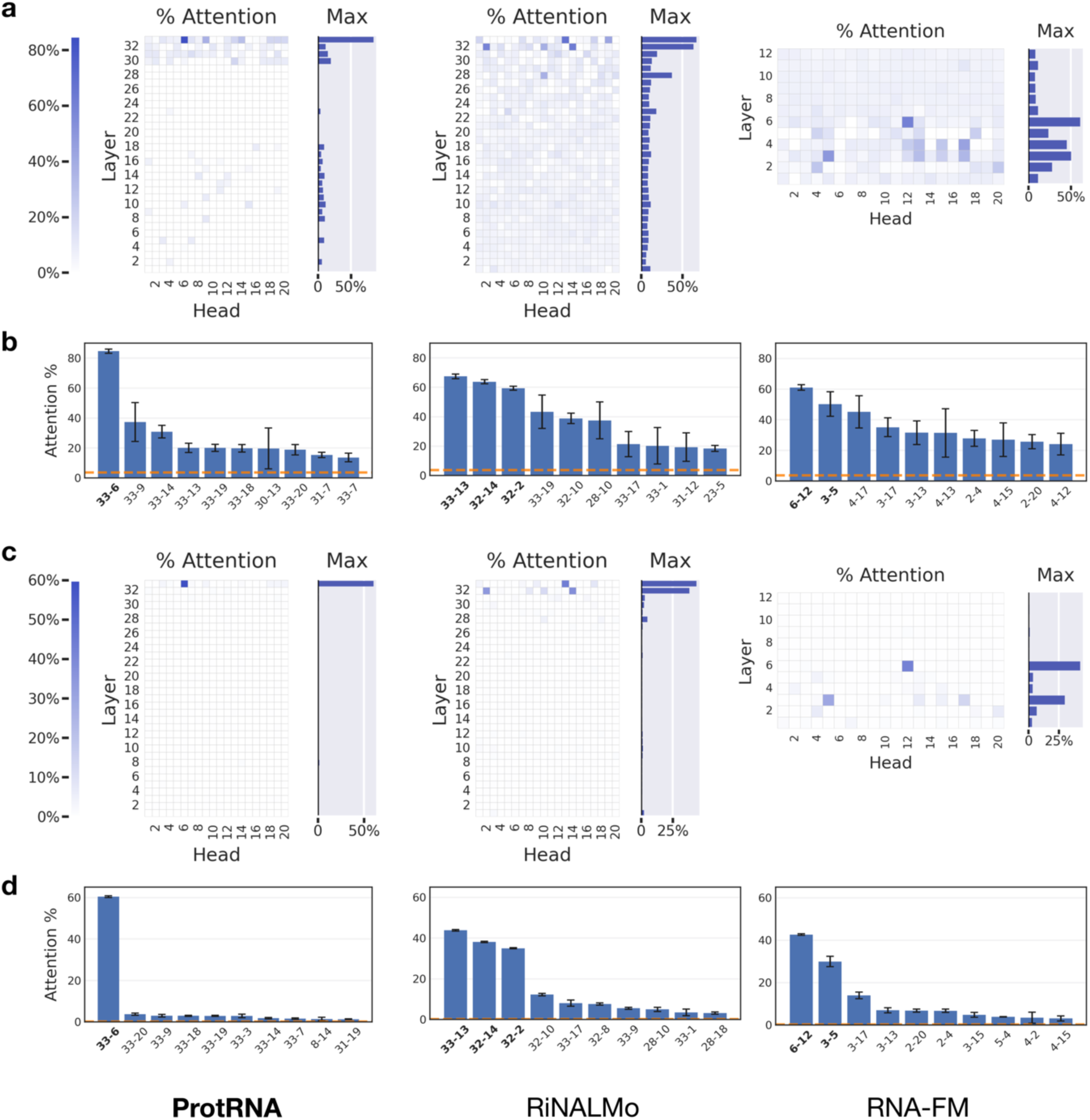
Attention analysis of ProtRNA against two baseline RNA language models. a-b, the results on contact map property. c-d, the results on secondary structure property. The ‘Attention%’ score for each head measures the proportion of its high attention weights that align with pairs in close contact among sequences from the RNA contact map dataset or which form hydrogen bonds across sequences from the RNA secondary structure dataset. The bar plots b and d under the heatmaps show the top 10 heads in the model with the highest scores, where the orange dashed lines represent the background frequency of the property under examination, and the black lines denote the confidence interval of the estimated scores under significance 0.05 with Bonferroni correction for multiple hypotheses.

As shown in the plot, ProtRNA exhibits a distinctive attention head pattern compared with the two other general RNA language models. Notably, ProtRNA features a highly specialized attention head, head 33-6, for both contact map and secondary structure properties. This head achieves a remarkable score of 84.6% for the contact map property, indicating that 84.6% of the high attention weights in this head correspond to pairs in contact. In contrast, RiNALMo and RNA-FM, while also employing shared structural heads for the contact map and secondary structure properties, depend on multiple heads with relatively lower scores. Specifically, RiNALMo mainly utilizes three such heads and RNA-FM employs two. This difference might stem from our constrained transfer learning strategy, where the ProtRNA model is required to compress information efficiently within just four unfrozen layers during pretraining on RNA sequence data.

### Evaluation of ProtRNA Representations on Downstream Tasks

#### Secondary Structure Prediction Using an Adapted Protein Structure Prediction Head

The secondary structure of an RNA molecule is the set of hydrogen-bond nucleotide base pairs formed within its primary sequence, including the Watson-Crick pairs A-U, G-C, and the Wobble pair G-U. Such base-pairing interactions induce secondary substructure motifs like stackings, loops, and bulges. Among the four levels of hierarchy in RNA structure, the secondary structure reveals basic information about the molecule’s function, characterizes the homology family it belongs to [30] [31], and facilitates construct design for tertiary structure determination [22].

Among all the deep learning models trained via direct supervised learning on secondary structure prediction, the following three models showed comparatively good results consistent across reports from different sources. SPOT-RNA deploys an ensemble of two-dimensional deep neural networks combining Residual Networks (ResNets) and Bidirectional Long Short-Term Memory cells (2D-BLSTMs) to output a 2D base-pair contact probability matrix [32]. The current SOTA DNN model UFold implements a U-Net architecture widely used in image segmentation tasks, to directly obtain a 2D contact map as SPOT-RNA, and applies post-process techniques to refine the results [33]. MXFold2 integrates deep learning with thermodynamic methods by folding in Zuker’s style to maximize the sum of Turner’s nearest neighbor free-energy parameters and folding scores computed from a deep neural network with convolutional layers and BLSTMs [34].

Another DNN-based methodology leverages transformer language models for sequence representation, similar to their application in protein structure prediction. A common practice in previous language models for secondary structure prediction involves training a relatively simple prediction head, such as the two-layer ResNet bottleneck used in RiNALMo. This approach exploits the self-attention mechanism, which constructs pairwise interactions between all positions in the sequence. The inductive bias inherent in this architecture effectively represents base pair interactions, facilitating accurate secondary structure predictions.

A new prediction head is proposed (Figure 1**.b**), which is an adapted AlphaFold2-style RotaFormer module from OPUS-Rota5 [35] and was originally used for highly accurate protein contact prediction. This advanced architecture has been shown to be more effective in extracting the embedded structural information in the language model representations. RNAErnie and RiNALMo also proposed their secondary structure prediction heads. The RNAErnie head stacks K downstream modules with shared parameters (referred to as the ‘STACK’ architecture in RNAErnie paper) to compute folding scores for each pair of bases. It then uses a Zuker-style dynamic programming approach to obtain the most favorable predictions. The RiNALMo head is a simple 2D bottleneck ResNet.

The performances of domain-specific models and pretrained language models with different prediction heads are evaluated on the bpRNA-1m [36] dataset under a unified pipeline. To compare the effectiveness of the RNA language model representations in this task, the raw sequence representations from the pretrained neural networks are utilized, with no fine-tuning or training dropout of the base model, as stated before. For the RNA language models, RNAErnie and RiNALMo were first tested with their own secondary structure prediction head and were then switched to the adapted RotaFormer head. Following the train-val-test split proposed in SPOT-RNA, the language model heads are all trained on TR0 and tested on TS0 of the bpRNA-1m dataset. For the RotaFormer head, the threshold is selected on VL0. For the domain-specific DNNs, we tested on the released models with training set as close to TR0 as possible. The post process techniques are left to the original architectural design of these models.

From **Table 2**, it is evident that our model outperforms others on TS0 for secondary structure prediction task. This highlights the capability of ProtRNA in predicting RNA secondary structures in the intra-family context even with significantly reduced trainable parameters and required training data of the pretrained language model. Additionally, our AF2-style RotaFormer head surpasses the RNAErnie or RiNALMo secondary structure prediction head in leveraging language model features. The advanced design of the RotaFormer head enables better utilization of the sequence representations, enhancing prediction accuracy. This demonstrates the superiority of our approach in integrating and interpreting sequence features. **Figure S2** demonstrates an example where the ProtRNA pretrained model with RotaFormer head predicts RNA secondary structure closer to the ground truth than the other two language models. The graph view in **Figure S2** is generated by RNAStructure [37].

**Table 2.** RNA secondary structure prediction performance of domain-specific models and language models on bpRNA TS0. RF stands for RotaFormer, the proposed secondary structure prediction head adapted from OPUS-Rota5. MCC refers to the Mathews Correlation Coefficient, Weisfeiler Lehman is proposed by RnaBench to compare structure graphs, and F1-shifted refers to the F1 score obtained after allowing flexible pairing.

| TS0 | F1 | MCC | Weisfeiler<br>Lehman | Recall | Precision | F1-shifted |
| --- | --- | --- | --- | --- | --- | --- |
| **Domain-Specific DNN** |  |  |  |  |  |  |
| SPOT-RNA | 0.587 | 0.596 | 0.725 | 0.673 | 0.545 | 0.624 |
| MXfold2 | 0.551 | 0.558 | 0.710 | 0.627 | 0.518 | 0.596 |
| UFold | 0.631 | 0.636 | 0.775 | 0.675 | 0.612 | 0.688 |
| UFold (from scratch) | 0.587 | 0.592 | 0.743 | 0.640 | 0.561 | 0.644 |
| **Language Model** |  |  |  |  |  |  |
| RNAErnie | 0.497 | 0.501 | 0.704 | 0.525 | 0.496 | 0.535 |
| RNAErnie-RF | 0.562 | 0.567 | 0.747 | 0.579 | 0.573 | 0.601 |
| RNA-FM | 0.528 | 0.540 | 0.757 | 0.536 | 0.579 | 0.565 |
| RNA-FM-RF | 0.624 | 0.628 | 0.789 | 0.622 | 0.647 | 0.670 |
| RiNALMo | 0.594 | 0.599 | 0.781 | 0.588 | 0.627 | 0.638 |
| RiNALMo-RF | 0.622 | 0.626 | 0.790 | 0.624 | 0.643 | 0.667 |
| ProtRNA | 0.646 | 0.651 | 0.807 | 0.644 | 0.672 | 0.694 |

In the secondary structure prediction task, the ProtRNA model consistently outperforms than the domain-specific DNNs and other language models on intra-family predictions, regardless of whether sequence homology is considered. This superiority is evident in bpRNA task (**Table 2**) and RnaBench [38] intra-family benchmark (**Table S2**). For inter-family structure prediction (**Table S3**), ProtRNA surpasses other language models, though it falls short compared to specialized deep learning models. However, the latter comparison with the domain-specific DNNs should be treated with conservation, as only inference on the test sets was conducted for these models. Additionally, we introduced experiments where UFold was trained from scratch on all three secondary structure prediction datasets. The corresponding results, labeled as “UFold (from scratch),” are presented in the tables.

Beyond canonical base pairs in secondary structure, RNA tertiary structures exhibit additional intramolecular interactions, including pseudoknots, non-canonical base pairs (not A-U, G-C, and G-U) and base-multiplets [39, 40][22]. Pseudoknotted base pairs are defined as the two base pairs (i, j) and (i’, j’) where i<i’<j<j’, forming non-nested pairs that are often excluded from standard dynamic programming algorithms for RNA structure prediction [22]. To assess the model’s ability to predict these tertiary interactions, we analyzed the performance of RNA language model heads—trained and validated on the RnaBench intra-family and inter-family tasks—alongside the domain-specific DNNs on these tertiary pairs. The evaluation was conducted on the benchmark datasets for these tasks.

The tertiary structure analysis results are presented in **Table S4** and **S5**, where the obtained results are similar to the secondary structure prediction case. For the intra-family task, the ProtRNA model outperforms all the other models in predicting pairs across all three types of tertiary contacts on the test set. For the inter-family task, ProtRNA performs comparably to RNA-FM on the inter-family task but lags behind RiNALMo. This limitation would be further discussed in **Discussion**.

The secondary structure prediction performance of ProtRNA across different RNA families is presented in Figure 4**.a** and **b**. F1 scores measuring the performance of the ProtRNA prediction heads trained on three standard datasets are shown. The models trained on intra-family and inter-family datasets, which have training sets five times larger than the bpRNA training set, exhibit similar performance across most RNA families. In contrast, the bpRNA model generally performs lower, particularly for families like telomerase.

**Figure 4.**
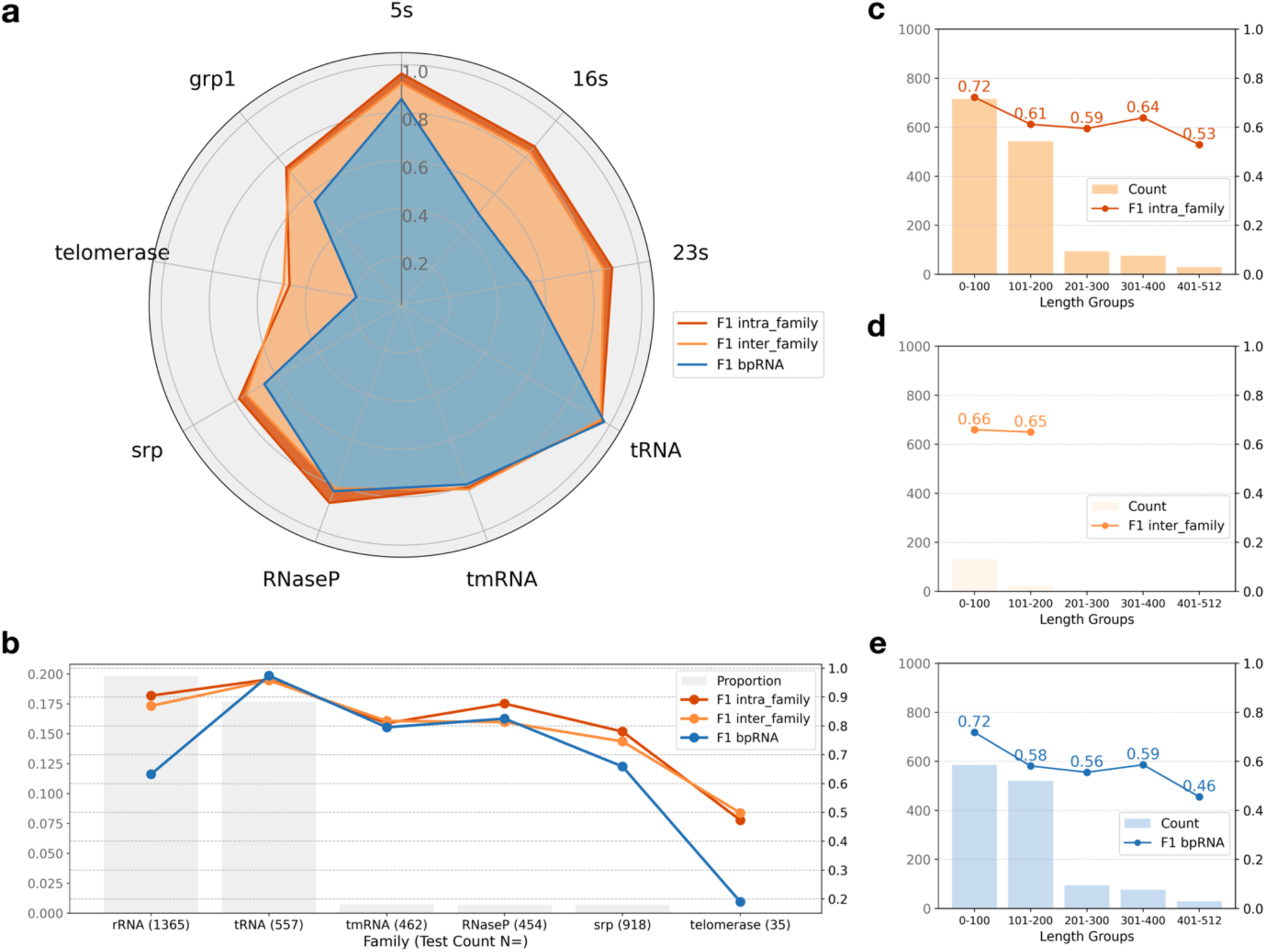
RNA secondary structure prediction performance of ProtRNA on different RNA families and across different sequence lengths. **a**, F1 scores measures of RNA secondary structure prediction performance on RNA sequences from different ncRNA families in ArchiveII, using models trained on intra-family, inter-family and bpRNA datasets respectively. **b**, prediction performance relative to the proportion of RNA families in the 6M pretraining data used by ProtRNA. The number in the parentheses after each family name is the test count of sequences in this family in the test set. The left vertical axis (in grey) represents the proportion of RNA sequences for each family in the pretraining data, while the right vertical axis shows the corresponding F1 scores. **c-e**, F1 scores of ProtRNA’s secondary structure prediction models (intra_family, inter_family, and bpRNA) across RNA sequence length groups. The left vertical axis represents sequence counts, while the right vertical axis shows the F1 scores.

All three models exhibit the lowest performance on the telomerase family, and the highest performance on 5s rRNA and tRNA families. Figure 4**.b** shows that this trend in F1 scores corresponds with the proportion of RNA families in the pretraining data. The rRNA family and tRNA family, which constitute 19.8% and 17.7% of the pretraining data, show better performance, while the telomerase family, which makes up only 0.01%, shows significantly lower performance.

The performance of three ProtRNA secondary structure prediction models across different sequence length groups is shown in Figure 4**. c-e**. All models exhibit a consistent decrease in F1 scores as the RNA sequence length increases. This trend likely reflects the limitations in performance as sequence length grows. The maximum sequence length of 512 represents the context length limit during pretraining, which may affect the models’ performance on sequences at or near this length.

#### Protein-RNA Interaction

The sequence of a biomolecule dictates its cellular function metaphorically to how code specifies a program, where the sequence acts as the code, which is ’compiled’ into a unique structure through folding, allowing the biomolecule to interact with other molecules and execute its function within the cell [41]. Thereby it is of interest how the sequence representations, yielded by compiling the sequence code to an extent through our transfer-learned language model and are showed above to contain structure information, could further facilitate tasks involving RNA interactions with other molecules in the cell.

Predicting RNA-binding protein (RBP) binding sites involves identifying specific regions on RNA molecules where RBPs interact. These interactions are crucial for various cellular processes, including RNA splicing, transport, stability, and translation [42] [43] . The task is complex due to the dynamic and context-specific nature of RNA structures and RBP interactions. Previous domain-specific DNN method, PrismNet [44], addresses the limitations of traditional methods by integrating deep learning with in vivo RNA structural data obtained using in vivo click Selective 2′-Hydroxyl Acylation and Profiling Experiment (icSHAPE), which allows the model to capture dynamic and condition-specific interactions between RBPs and RNA.

Here we reproduced PrismNet but substituted the icSHAPE experimental RNA structural data with RNA sequence representations obtained from our ProtRNA model and other benchmarking pretrained RNA language models (see Figure 1**.c**). Using the HeLa cell dataset, we aim to test if our ProtRNA sequence representations could effectively facilitate the RBP prediction task.

As shown in Figure 5, compared with the baseline seq-only model, ProtRNA + Seq consistently demonstrates positive impacts and only minor negative impacts in most cases. This performance matches or surpasses that of the RealSS + Seq model (see Figure 5**.a**), which utilizes icSHAPE RNA structural data. ProtRNA + Seq also outperforms the other three models (see Figure 5**.b-d**): RNA-FM and RNAErnie exhibit notable negative impacts for many proteins, and RiNALMo largely fails to achieve the improvements seen with ProtRNA or RealSS. The overview of the performance comparison of RealSS and four language models could be found in **Figure S2**. The performance on the protein-RNA interaction task suggests that, despite using less training data on RNA sequences and fewer trainable parameters, our pretraining strategy enhances the model’s ability to handle the complex interaction task between proteins and RNA. This may be attributed to the pretraining process that incorporates biomolecules of different modalities.

**Figure 5.**
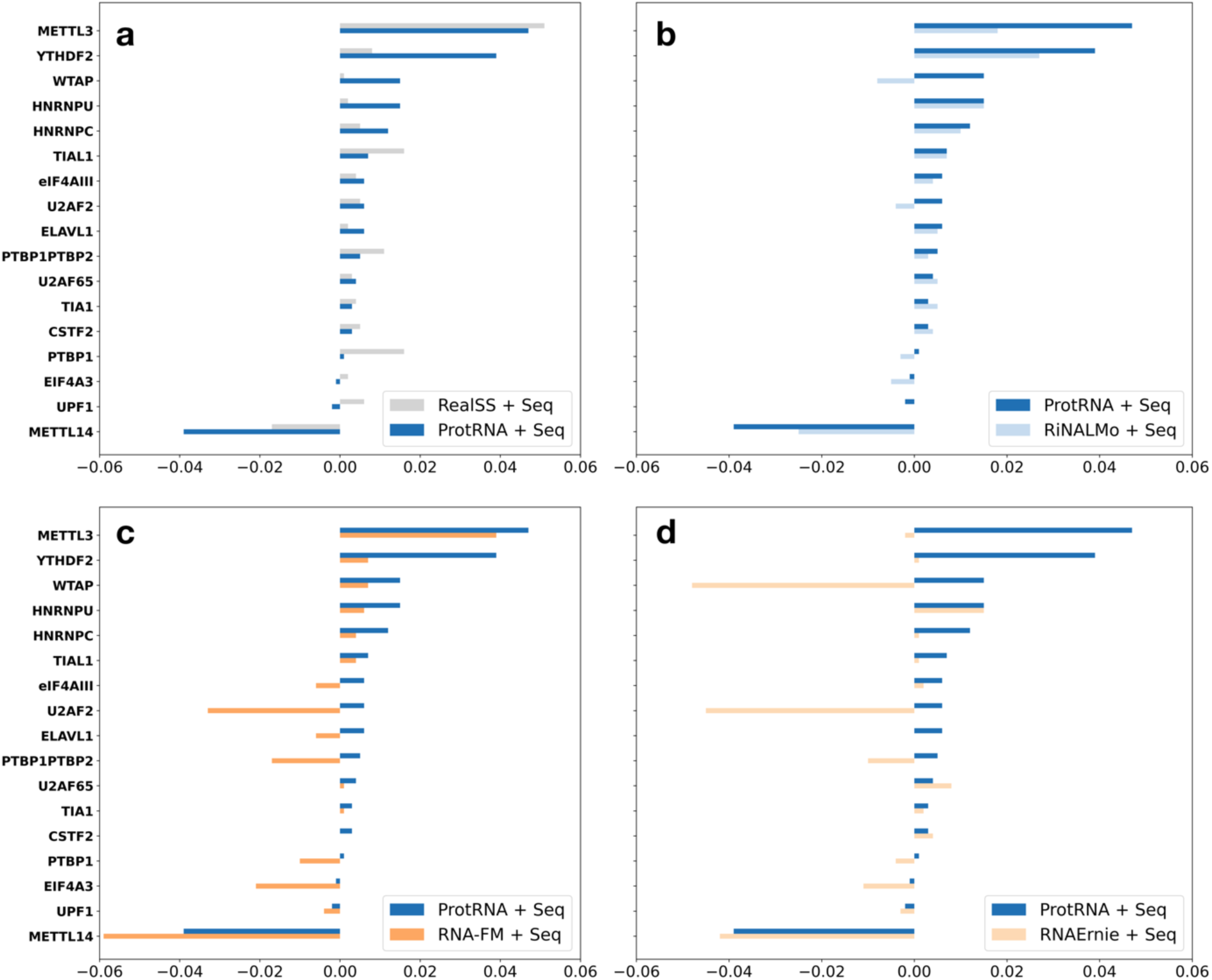
Pairwise Comparison of Protein-RNA interaction performance (AUROC) between RNA language model features on 17 RBPs in Hela Cell. “Seq” stands for one-hot representations of the RNA sequence. “RealSS” refers to the 1-dimensional experimental icSHAPE data of the RNA secondary structure. “+” means the features are concatenated as inputs to the PrismNet model. The y-axes are the 17 RBPs, and the x-axes are the area under the receiver operating characteristic (AUROC) score of each model. The AUROC scores are shifted to compare with the model performance with inputs of “Seq” only.

To evaluate the performance of each PrismNet head, we generate a Receiver Operating Characteristic (ROC) curve by thresholding the predicted scores to derive the corresponding true positive and false positive rates. For each threshold, predictions with scores greater than or equal to the threshold are classified as positive, and those below as negative. This procedure produces a series of True Positive Rate (TPR) and False Positive Rate (FPR) pairs that are used to construct the ROC curve. The Area Under Curve (AUC), shown in Figure 5, is then computed from these TPR and the FPR pairs to provide a compact scalar summary of model performance. For each model, **Table S6** presents the average TPR, FPR, and their difference across different RBPs in Hela cells. Additionally, **Figure S4** shows the confusion matrices for different model representations, generated at the threshold that maximizes the difference between TPR and FPR. See the **Methods** section for further details on these metrics.

#### Mean Ribosome Loading

Mean Ribosome Loading (MRL) refers to the average number of ribosomes bound to a single mRNA molecule during translation. It measures translation efficiency, indicating how actively a particular mRNA is being translated into protein. The 5’ untranslated region (5’ UTR) of mRNA regulates protein translation efficiency, as it contains specific sequences and structural motifs crucial for the binding of ribosomes and translation initiation factors [45] [46]. Accurate prediction of MRL from the 5’UTR of an mRNA sequence would significantly enhance drug design in mRNA therapeutics.

Predicting MRL from 5’UTR is a regression task that involves creating models capable of capturing the regulatory features within the 5’UTR and predicting their impact on ribosome loading. Advanced machine learning techniques, particularly deep learning models, are employed to learn these complex relationships from large datasets of 5′ UTR sequences and their corresponding MRL values [47] [48].

In this task, we aim to test our model’s generalizability to the structural information contained in the 5’UTR of mRNA sequences, which is a noncoding region that serves a function role. The training dataset and the evaluation datasets, Human7600 and Random7600 are provided by [49]. We utilized the same prediction head architecture as in Uni-RNA and RiNALMo, featuring a simple 1D ResNet (see Figure 1**.d**). For all RNA language models under comparison, the sequence representations are from pretrained models, and only the prediction head is trained. During training, R^2^ is used as the evaluation metric, as in RiNALMo.

As seen from **Table S7**, our ProtRNA representations of the 5’UTR achieve higher R^2^ and lower Mean Absolute Error (MAE) in the regression task. This indicates the better generalization ability of ProtRNA model to RNA sequences out of distribution of the pretraining data, compared with the other RNA language models that are also trained solely on ncRNA data. These results underscore the generalizability of our ProtRNA representations.

## Discussion

In this paper, we introduced a cross-modality transfer learning strategy for RNA language model pretraining, leveraging model parameters from ESM-2, which is extensively trained on the protein language, a ‘resource-rich’ modality. Our ProtRNA model, built on this transfer learning approach, achieves comparable or even superior performance in several RNA downstream tasks when compared to other RNA foundation models and domain-specific deep learning models. Such performance highlights that the effectiveness of transferring information from a pretrained protein language model to RNA sequences.

Specifically, a new secondary structure prediction architecture is proposed, shown in Figure 1**.b**, which combines language model representations with an advanced AF2-style RotaFormer module. This architecture demonstrates superior performance (**Table 2**), adding up to the DNN-based toolbox in RNA secondary structure prediction. The performance of ProtRNA representations in the protein-RNA interaction task (Figure 5) highlights the effectiveness of the encoded information, resulting from evolutional transfer learning pretraining, to facilitate tasks involving the interaction between biomolecules of different modalities. Finally, ProtRNA generalizes to produce mRNA 5’UTR representations that achieve better MRL prediction accuracy (**Table S7**).

Additionally, since the ProtRNA model achieves competitive performance in RNA-specific tasks while retaining most of the transformer layer parameters from the protein model ESM-2, it may serve as an intriguing subject for future studies on the interpretability of deep learning models within a biological context. Two preliminary analyses were conducted: the t-SNE plot demonstrates the ProtRNA representation space with clearly segregated protein and RNA sequences; the attention analysis reveals a distinctive attention head pattern of the ProtRNA model related to RNA structural properties. While further analysis to understand the mechanisms behind this performance is beyond the scope of this work, it is a promising avenue for future research that could yield insights into the interplay between different biological modalities and inform the development of improved models.

Despite the success of the ProtRNA transfer learning strategy, as demonstrated by its performance across various downstream tasks, some limitations remain. First, the biased distribution of RNA families in the pretraining data influences the model’s secondary structure prediction performance. The model exhibits a bias toward families with higher representation, such as 5s rRNA and tRNA (which comprise 15%-20% of the pretraining dataset), while achieving the lowest performance on the telomerase family, which accounts for 0.01% of the pretraining data. In the secondary structure prediction for challenging base pairs derived from tertiary structures (**Table S4** and **S5)**, ProtRNA performs comparably to RNA-FM on the inter-family task but lags behind RiNALMo. This discrepancy might have also arisen from the differences in pretraining dataset: ProtRNA and RNA-FM are both trained exclusively on 23M sequences from RNAcentral, whereas RiNALMo incorporates additional databases, yielding a larger and more diverse 36M pretraining dataset. The potential bias introduced by the overrepresentation of certain RNA families in RNAcentral has also been noted in recent studies [9]. Future work incorporating a more comprehensive and diverse pretraining dataset could help mitigate this limitation. Second, as shown in **Table S8**, the ProtRNA model exhibits significant catastrophic forgetting when applied to protein downstream tasks. This suggests that while the model effectively captures RNA structural and functional features, its ability to retain previously learned protein-related knowledge diminishes. Addressing this challenge may require strategies such as continual learning or multi-modal training to better preserve cross-domain knowledge.

For our proposed cross-modality transfer learning strategy, the current model only utilizes data from the target modality, in our case, RNA. To optimize the model performance on the target modality, other established methods in cross-lingual transfer learning may also be adopted, such as the representation alignment or joint training objectives between two languages [50] [51]. However, it is crucial to note that the mapping relationship between protein and ncRNA sequences differs significantly from that in natural languages. Therefore, adapting these NLP methods requires careful consideration. For instance, when constructing training batches with sequence pairs of ncRNAs and proteins to analogously match their “meaning”, one might consider incorporating structural or functional motifs. In contrast, for mRNA, the mapping between proteins and mRNA follows a straightforward codon rule, which could be captured easily by a well-trained neural network. Thus, alignment training between these two modalities may not add significant semantic value. Taking the above considerations into account, more delicate cross-modality training strategies suitable for protein and RNA, or even other modalities, could be explored in the future. Additionally, unfreezing more layers during pretraining or fine-tuning the full model for downstream tasks might further improve performance.

## Methods

### ProtRNA Pretrained Language Model

#### Pretraining Data

The original dataset, collected from the latest RNAcentral release when the experiments are conducted, contains 38 million RNA sequences. To reduce sequence redundancy, we applied MMSeqs2 algorithm with parameters set to form clusters with 90% sequence identity and representative sequences covering at least 95% of the cluster members. This process reduced the dataset to 6 million representative sequences, resulting in an 84% reduced sparse dataset. This reduction is comparable to the UniRef50 dataset used for ESM-2 pretraining, which achieved a 79% reduction from the original UniRef100 dataset.

#### Tokenizer

Similar to other general ncRNA language models, RNA sequences in ProtRNA are tokenized at single-nucleotide resolution, treating each nucleotide base as a separate token. The vocabulary expands upon the original ESM-2 amino acid tokens by including nucleotide tokens (“a”, “c”, “u”, “g”, and “x”). Lowercasing distinguishes nucleotide tokens from uppercase amino acid tokens, and the “x” token represents other non-common modified or unknown nucleotides. Special tokens like <CLS>, <PAD>, <EOS>, and <MASK> utilize the same encoding and embeddings as in ESM-2 to leverage learned embeddings.

#### Transfer Learning Pretraining Method

For transfer learning pretraining, we leveraged fixed parameters from the 650M parameters ESM-2 model, with 33 transformer blocks and 1280-embedding dimension, that was sufficiently pretrained on protein data UniRef50/D. We applied minimum modifications to the original model architecture to fully utilize the evolutional information stored in its attention head weights. Namely, we only extended the embedding layer and the bias layer to accommodate the size of added RNA tokens. During training, most of the transformer blocks in the protein model ESM2, i.e., 29 among 33 (≈88%) of the blocks, were fixed.

We allow a context window of 512. Our model is pretrained on 4 V100 GPUs of 32GB memory for 6 epochs. During 5000 warm-up steps, we gradually increase the learning rate to 1e-4 and use Adam optimizer, and then Nadam optimizer is applied with the constant learning rate 1e-4. The pretraining task for our architecture is as suggested in RoBERTa [52], where masked language modeling is applied with 15% tokens set to be targets and replaced with 80% the time a <MASK> token, 10% a random token from the vocabulary, and 10% the original token for robust learning. The loss function is the classic cross-entropy loss for multi-class prediction. And the weight updates are based on the average loss per batch.

### Attention Analysis

#### Metric

The metric 𝑝_α_(𝑓) measures the alignment of attention weights with underlying properties of token pairs, in our case, contact map or secondary structure base pairing. Indicator function 𝑓(𝑖, 𝑗) returns 1 if the property is present in token pair (𝑖, 𝑗), 0 otherwise. 𝜃 is the threshold for high attention weights. For each attention head, the score is aggregated for all sequences 𝑥 over the dataset 𝑋. In our analysis, the threshold 𝜃 is set to be 0.3, following the practice in [4].

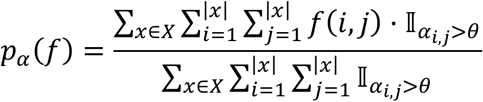

#### Dataset

The contact map dataset is the whole dataset as provided in the RNA benchmark BEACON [53], while the secondary structure dataset is composed of 5000 sequences randomly chosen from the training set TR0 of the bpRNA dataset.

### Baseline RNA Language Models for Downstream Evaluation

In this paper, we chose RNA-FM, RiNALMo, and RNAErnie as baseline RNA language models, which are open-resource models with reported competent performance. When evaluating the model performance on downstream tasks, we have restrained from fine-tuning the base model or employing typical training techniques such as dropout. These restraints ensure a fair comparison among foundational RNA language models, thereby providing a clearer indication of the quality of the extracted features.

### Secondary Structure Prediction

#### Dataset

bpRNA-1m [36] is a benchmark in RNA secondary structure prediction for deep learning methods, with the training, validation and test split proposed in SPOT-RNA [32]. It contains over 100, 000 structures collected from seven sources, including consensus structures derived from comparative sequence analysis and also X-ray crystallography and NMR techniques from Protein Data Bank (PDB).

In addition to the widely used bpRNA-1m dataset, we utilized two datasets provided by the RnaBench pipeline which removes sequence and structure homologs between the training, validation and test partition of the datasets. Specifically, the intra-family benchmark accounts for sequence homology, and the inter-family benchmark additionally considers structure homology.

#### RotaFormer Head

As illustrated in **Figure 1.b**, the RotaFormer head adopts the backbone architecture of the RotaFormer module for protein structure prediction with several modifications. The adapted head utilizes both 1D and 2D features as inputs and deploys 12 blocks of Sequence to Sequence (Seq2Seq) submodules and Pair to Pair (Pair2Pair) submodules. The Seq2Seq submodules feature Sequence Row Attention with Pair Bias, while the Pair2Pair submodules feature Triangle Multiplication and Triangle Attention. This is followed by a 5x5 kernel CNN to obtain the final probability map for base-pairing.

#### Benchmarking Domain-Specific DNNs

For the secondary structure domain-specific deep learning models, we obtained their released weights and tested their performance on TS0, intra-family, and inter-family datasets under a unified pipeline. In the following we refer to the original train-validation-test split data from bpRNA-1m as TR0, VL0, and TS0. The SPOT-RNA model was trained on TR0 and then fine-tuned on a curated small high-resolution structure data from PDB, which is the only weight available. MXFold2 was trained on TR0 and validated on VL0 from bpRNA-1m. And for UFold, we utilized its released ufold_train.pt weight. In addition, we trained UFold from scratch on all three tasks using the default training schedule provided in its codebase. The best-performing epoch was selected based on the highest F1 score achieved on the corresponding validation set, with early stopping applied using a patience of 15 epochs. Training from scratch for SPOT-RNA and MXFold2 on RnaBench was not included due to following concerns: SPOT-RNA lacks publicly available training modules and scripts for preparing training data; MXFold2 is limited to pseudoknot-free predictions, making it incompatible with the RnaBench dataset, which contains a non-negligible number of pseudoknotted structures critical for evaluation.

#### Post-Process

Typically, deep-learning methods output a 2D contact interaction map, which for certain models necessitates additional post-processing techniques to derive definitive base-pairs from the probability map. These methods often involve enforcing symmetry on the contact matrix. For instance, UFold screens out non-canonical base pairings and restricts sharp loops. In contrast, the RiNALMo-head leverages a greedy algorithm to iteratively establish base pairs with the highest pairing probabilities while excluding potential clashing pairs.

#### Training and Evaluation

During the training of the RotaFormer head, the Adam optimizer was used with an initial learning rate of 2e-4, which was halved every five epochs. The head was trained on 4 NVIDIA Tesla V100 GPUs with a total batch size of 4. For each head trained with language model representations, the best epoch was selected based on the highest F1 score at the optimal threshold, using early stopping with a patience of 3 epochs. The selected best models then produce base pairs predictions with their respective post-process. The metrics from the RnaBench pipeline were applied to these predicted base pairs to obtain the results in **Table 2**, **Table S2-5**.

#### Metrics

It’s worth noting that, as conformational dynamics exist in base pairing, which means that base pairs can fluctuate, the (i±1, j), (i, j±1) alternative states regarding the (i, j) base pair should also be considered correct in secondary structural prediction [22]. This is often reflected in the metric F1-shifted in secondary structure prediction task. Different methods were evaluated by obtaining each method’s predicted base pairs and applying the metrics provided in the RnaBench pipeline.

### Protein-RNA Interaction

#### Dataset

From the Hela clip data provided by PrismNet, for Seq+SS model, the icSHAPE data are utilized, while for other models, only the RNA fragments sequences are taken as input. According to the original setting in PrismNet, 20% of the samples are taken as test set. Differently, we added train-validation split that randomly takes out 20% of the training set as validation set. The random seed is set the same across different models as default.

#### Training and Evaluation

The PrismNet heads were trained using default hyperparameters: a learning rate of 0.001, batch size of 64, and a linear scheduler warmup with a weight decay of parameter 1e-06. Each model was trained on a single NVIDIA Tesla V100 GPU. The best epoch for each model was selected based on the highest epoch validation AUC, with early stopping applied and a patience of 20 epochs. The selected best models were then evaluated on a separate test set to obtain the final AUC values presented in **Figure 5**.

#### Metrics

To evaluate the performance of each PrismNet head, we generate a Receiver Operating Characteristic (ROC) curve using the *roc_curve* function from the scikit-learn library. This function takes the predicted scores and their corresponding ground-truth labels as inputs. It sorts the unique predicted scores in descending order and treats them as potential decision thresholds. For each threshold, predictions with scores greater than or equal to the threshold are classified as positive, while the remainder are classified as negative. This procedure produces a series of True Positive Rate (TPR) and False Positive Rate (FPR) pairs used to construct the ROC curve. For each model, **Table S6** presents the average TPR, FPR, and their difference across different RBPs in Hela cells.

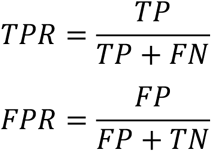

The Area Under Curve (AUC) values shown in **Figure 5** are computed using the *auc* function from the scikit-learn library applied to the TPR and FPR pairs. The AUC provides a compact scalar summary of the model’s performance. During validation, AUC is used to select the best-performing model, while test AUC values across different models underscore distinctions in language model representations.

In addition to the mean TPR and FPR values, we present the confusion matrices for the methods in **Figure S4**. For each PrismNet head, the optimal threshold is determined on the validation set by maximizing the difference between TPR and FPR:

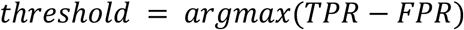

This selected threshold is then applied to the test set to compute the numbers of True Positives (TP), False Positives (FP), True Negatives (TN), and False Negatives (FN). The average TP, FP, TN, and FN values for each model are further weighted according to the number of test observations for each protein.

#### Feature Reduction

To optimize the utilization of language model features in PrismNet, we performed dimensionality reduction on the original high-dimensional features. Specifically, language features were reduced to 32 dimensions using a trainable linear layer. Similarly, one-hot sequence embeddings were transformed to 16 dimensions to maintain consistency in magnitude. Hyperparameters were aligned with PrismNet settings: the sequence-only model utilized a kernel height of 3, whereas models incorporating icSHAPE data or language model features utilized a kernel height of 5.

### Mean Ribosome Loading

#### Dataset

The dataset used is a large-scale synthetic Human 5′ UTR library comprising 5′ UTR sequences with measured MRL values originally described by [49]. It includes human and random UTR sequences of varying lengths, with 100 sequences per length (from 25 to 100 nucleotides) selected based on deep read coverage. This sampling also created two evaluation datasets, Random7600 and Human7600, ensuring balanced representation across sequence lengths.

#### Head Architecture

The MRL prediction head is the same as the one used by RiNALMo, despite for RNA-FM feature input, the feature dimension of the first linear layer is 640 instead of 1280 for ProtRNA and RiNALMo.

#### Training and Evaluation

During the training of the MRL prediction head, the Adam optimizer was used with an initial learning rate of 1e-4. A linear learning rate scheduler was applied, starting at 100% of the initial learning rate and decaying to 10% over the course of 5000 iterations. The prediction head was trained across 4 NVIDIA Tesla V100 GPUs, with a total batch size of 64. For each head trained with language model representations, the best epoch was selected based on the lowest Mean Absolute Error on the validation set, with early stopping applied and a patience of 3 epochs.

## Data availability

The code repository is available at https://github.com/roxie-zhang/ProtRNA. The pretrained model weights and datasets can be downloaded from https://zenodo.org/records/14795554.

## Acknowledgements

J.M. wants to thank the support from the National Key Research and Development Program of China (No. 2024YFA1307502), the Science and Technology Innovation Plan of Shanghai Science and Technology Commission (No. 23JS1400200), and the Research Fund for International Senior Scientists (No. W2431060). G.X. wants to thank the support from the National Natural Science Foundation of China (No. 32300535).

## Author Contributions

R.Z. and G.X. jointly developed and trained the model, with R.Z. writing the primary scripts and G.X. refining the implementation. R.Z. also performed the analyses and drafted the manuscript. R.Z., B.M., and G.X. collaborated on downstream experiments. J.M. and G.X. supervised the research and designed the study. All authors contributed to project discussions and manuscript revisions.

## Notes

https://github.com/roxie-zhang/ProtRNA

### Competing Interest Statement

The authors have declared no competing interest.

### Summary of Updates

Update on the section of secondary structure prediction to provide inter-family evaluation; Refinement of text, graphs and tables for clarity; Add ablation studies for the pretraining strategy; Supplemental files updated

