## supplemental materials for "ProtRNA: A Protein-derived RNA Language Model by Cross-Modality Transfer Learning"

Zhang et al.

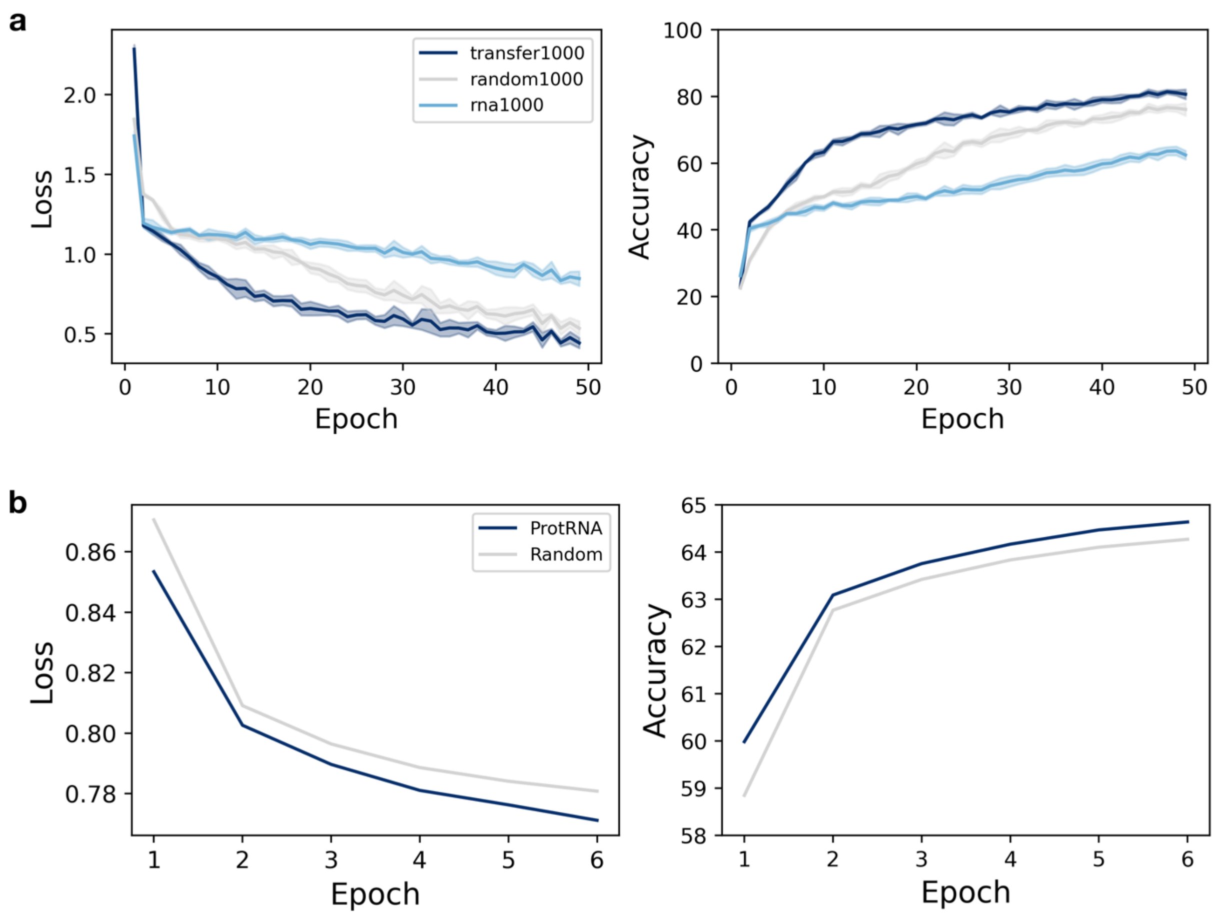


**Figure S1. Pretraining comparison between transfer learning method and training from scratch. a,** three models–transfer1000, rna1000, and random1000–were trained on a small dataset consisting of 1,000 randomly selected ncRNA sequences from RNAcentral. transfer1000 is initialized from the same checkpoint as the main ProtRNA model, with the last four transformer blocks unfrozen during training, whereas random1000 shares the same architecture and unfrozen layers but is initialized at random. In contrast, rna1000 comprises four transformer blocks with randomly initialized weights. All three models are designed to with an identical number of trainable parameters to focus the comparison on the impact of initialization and training strategy. The performance curves in both panels represent the mean values, with shaded regions indicating one standard deviation from five independent runs with different random seeds. **b,** the training trajectory of the ProtRNA model compared with Random, a model initialized at random, on full pretraining data. The loss metric for each epoch is computed as the sparse categorical cross-entropy between the true token indices and the model's predicted logits, averaged over all masked tokens across all batches in the epoch. The accuracy metric for each epoch is defined as the percentage of correct predictions in the Masked Language Modeling (MLM) pretraining task, averaged across all batches in the epoch.

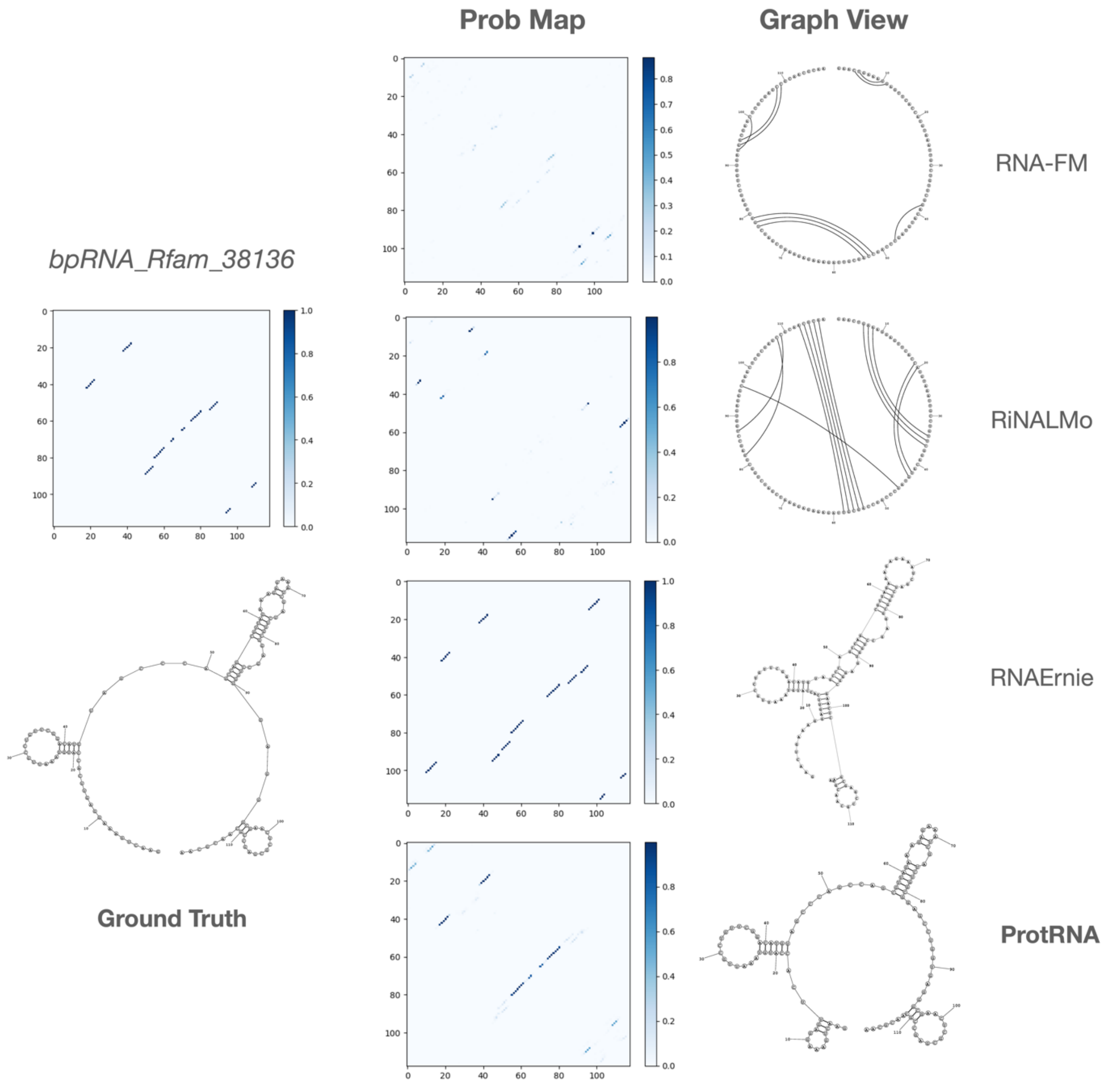


**Figure S2.** The secondary structure probability maps and binary pairing in graph view from the prediction of the three RNA language models on a sequence from bpRNA dataset. Note that due to the design of the RNAErnie head, the Prob Map displayed is the bool matrix base pairs predictions.

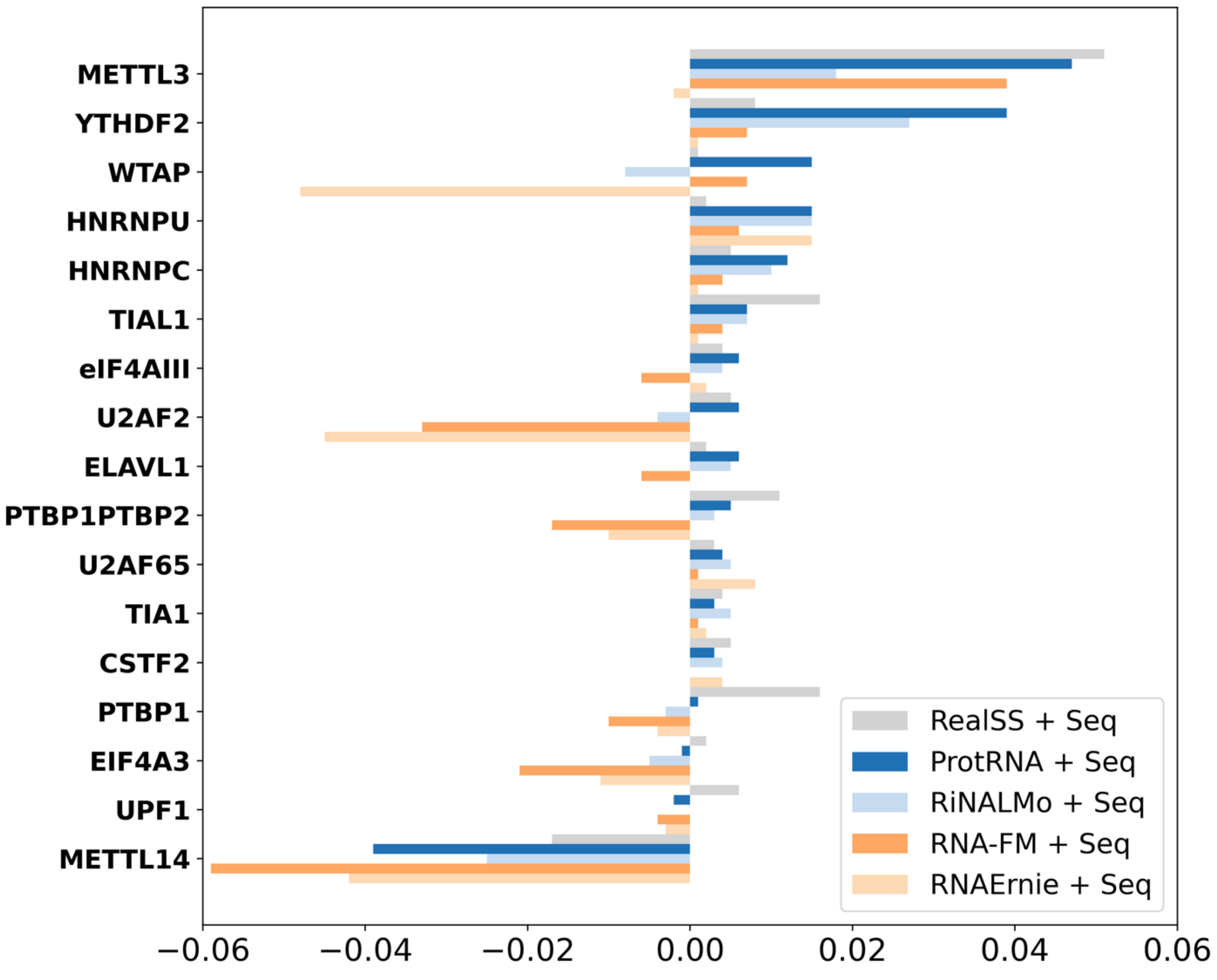


**Figure S3. Overview of Protein-RNA interaction performance (AUROC) of the RNA language model and comparative models on 17 RBPs in HeLa cells.** "Seq" stands for one-hot representations of the RNA sequence. "RealSS" refers to the 1-dimensional experimental icSHAPE data of the RNA secondary structure. "+" means the features are concatenated as inputs to the PrismNet model. The y-axes are the 17 RBPs, and the x-axes are the area under the receiver operating characteristic (AUROC) score of each model. The AUROC scores are shifted to compare with the model performance with inputs of "Seq" only.

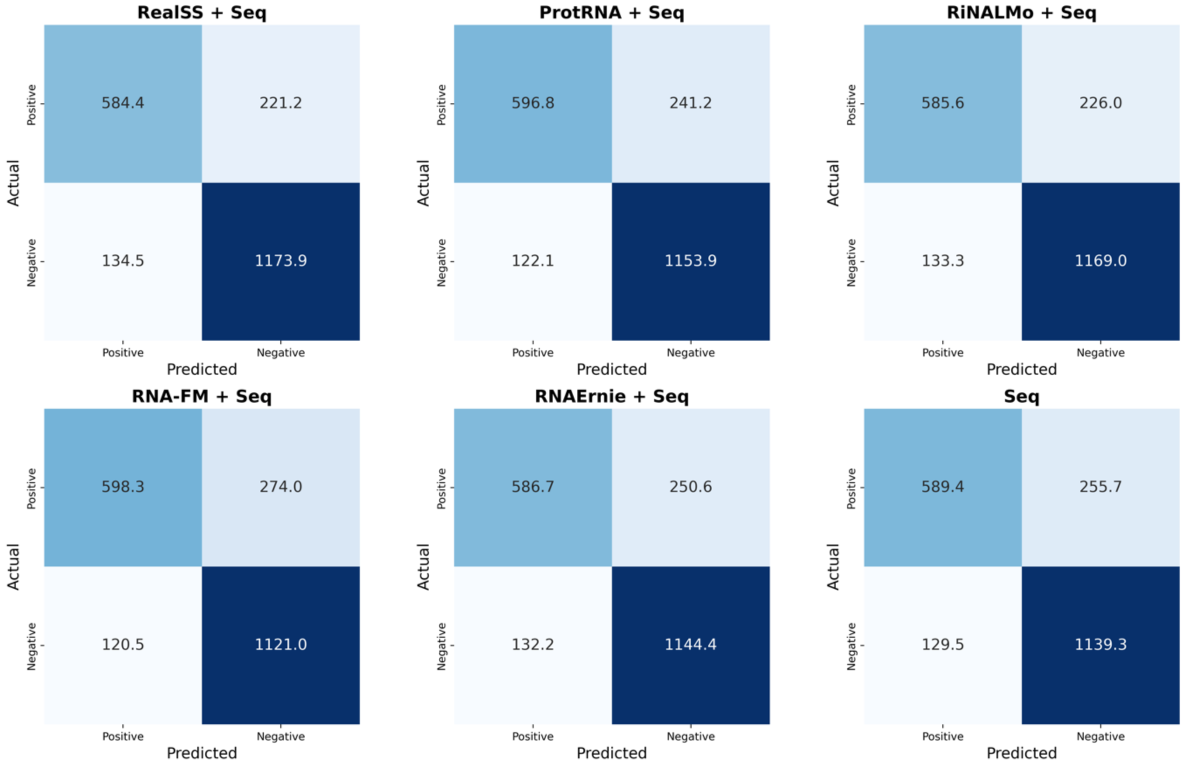


**Figure S4. Confusion matrices for each method across 17 RBPs in Hela cells.** Thresholds for each method under different proteins were selected by maximizing the difference between TPR and FPR on the validation set. Average TP, FP, FN, and TN values are weighted by the number of test observations for each protein.

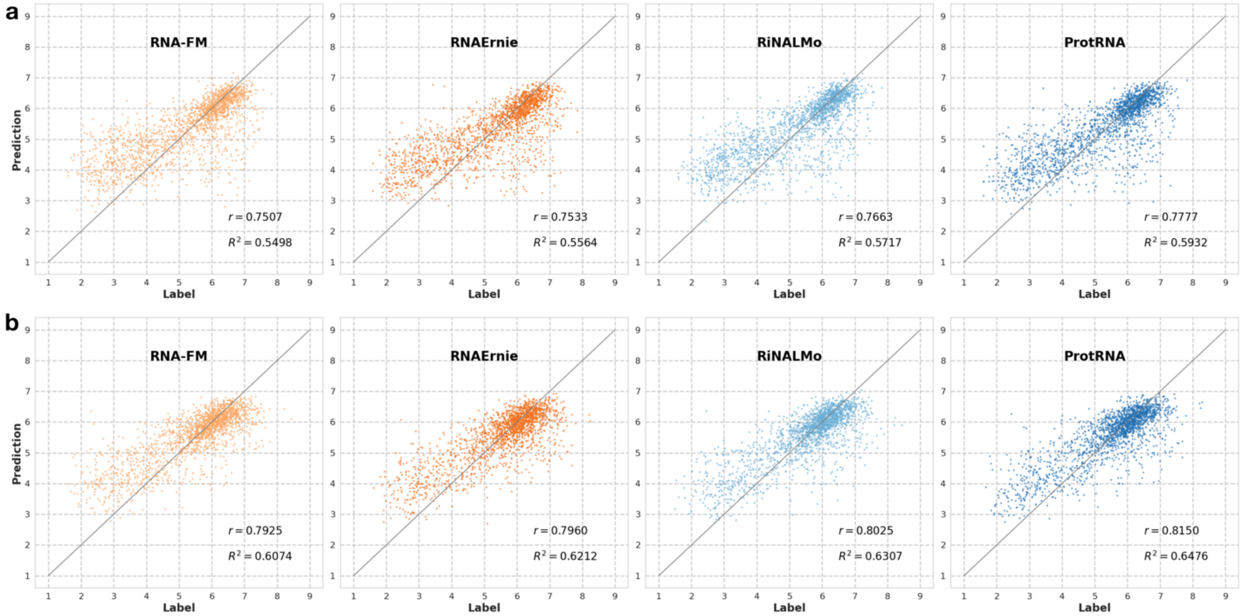


**Figure S5. Head-to-head comparison of language models mean ribosome loading prediction performance**. 'r' stands for Pearson's correlation coefficient. **a,** performance evaluated on random7600 dataset. **b,** performance evaluated on human7600 dataset.

**Table S1. Comparison of secondary structure prediction performance on bpRNA TS0 between models pretrained with different efficient transfer learning strategies. LoRA:** Low-Rank Adaptation following common practice for language models, with rank set to 16. **Random:** pretraining from randomly initialized starting weights of the ESM-2 architecture. **ProtRNA (L4)**: transfer learning with the last 4 unfrozen transformer layers from ESM-2 weights, following the same pretraining strategy as the main model. **L2**, **L6**: transfer learning with 2 or 6 unfrozen layers respectively from ESM-2 weights. **epoch** means pretraining epoch for the language model. The secondary structure prediction RotaFormer head is trained and validated in the same procedure as for the main model ProtRNA. Best F1 scores across models at each pretraining epoch is shown in bold. The best F1 for each epoch is shown in bold.

| **epoch/F1** | **LoRA** | **Random** | **ProtRNA (L4)** | **L2** | **L6** |
|---|---|---|---|---|---|
| **e1** | 0.611 | 0.614 | 0.626 | **0.632** | 0.623 |
| **e2** | 0.615 | 0.623 | 0.634 | 0.628 | **0.645** |
| **e3** | 0.614 | 0.631 | 0.637 | 0.627 | **0.644** |
| **e4** | 0.629 | 0.634 | 0.633 | 0.630 | **0.645** |
| **e5** | 0.620 | 0.632 | **0.646** | 0.628 | 0.643 |

**Table S2. RNA secondary structure prediction performance of domain-specific models and language models on RnaBench intra-family benchmark dataset.**

| Intra-Family Benchmark | **F1** | **MCC** | **Weisfeiler Lehman** | **Recall** | **Precision** | **F1-shifted** |
|---|---|---|---|---|---|---|
| **Domain-Specific DNN** | | | | | | |
| SPOT-RNA | 0.601 | 0.610 | 0.723 | 0.671 | 0.576 | 0.637 |
| MXfold2 | 0.565 | 0.573 | 0.709 | 0.623 | 0.552 | 0.609 |
| UFold | 0.633 | 0.638 | 0.765 | 0.661 | 0.635 | 0.687 |
| UFold (from scratch) | 0.628 | 0.634 | 0.763 | 0.658 | 0.628 | 0.686 |
| **Language Model** | | | | | | |
| RNA-FM | 0.581 | 0.591 | 0.758 | 0.593 | 0.622 | 0.617 |
| RiNALMo | 0.603 | 0.609 | 0.768 | 0.598 | 0.641 | 0.644 |
| ProtRNA | 0.665 | 0.670 | 0.792 | 0.676 | 0.680 | 0.711 |

**Table S3. RNA secondary structure prediction performance of domain-specific models and language models on RnaBench inter-family benchmark dataset.**

| Inter-Family Benchmark | **F1** | **MCC** | **Weisfeiler Lehman** | **Recall** | **Precision** | **F1-shifted** |
|---|---|---|---|---|---|---|
| **Domain-Specific DNN** | | | | | | |
| SPOT-RNA | 0.721 | 0.730 | 0.706 | 0.652 | 0.841 | 0.745 |
| MXfold2 | 0.686 | 0.699 | 0.704 | 0.592 | 0.846 | 0.714 |
| UFold | 0.644 | 0.661 | 0.673 | 0.543 | 0.831 | 0.674 |
| UFold (from scratch) | 0.670 | 0.677 | 0.693 | 0.602 | 0.784 | 0.704 |
| **Language Model** | | | | | | |
| RNA-FM | 0.584 | 0.594 | 0.628 | 0.508 | 0.725 | 0.616 |
| RiNALMo | 0.619 | 0.622 | 0.643 | 0.587 | 0.682 | 0.654 |
| ProtRNA | 0.658 | 0.668 | 0.672 | 0.585 | 0.788 | 0.687 |

**Table S4. Secondary structure prediction comparison on intra-family benchmark dataset for pseudoknots (pk), multiplets, and noncanonical pairs (nc).** '-' indicates that the model either does not support predicting these interactions by design or has this feature disabled by default.

| Intra-Family Benchmark | | **F1** | **MCC** | **Weisfeiler Lehman** | **Recall** | **Precision** | **F1-shifted** |
|---|---|---|---|---|---|---|---|
| **SPOT-RNA** | pk | 0.236 | 0.245 | 0.723 | 0.246 | 0.270 | 0.246 |
| | multiplet | 0.099 | 0.104 | 0.801 | 0.090 | 0.137 | 0.120 |
| | nc | 0.212 | 0.230 | 0.929 | 0.262 | 0.245 | 0.225 |
| **MXFold2** | pk | - | - | - | - | - | - |
| | multiplet | 0.0 | 0.0 | 0.0 | 0.0 | 0.0 | 0.0 |
| | nc | - | - | - | - | - | - |
| **UFold** | pk | 0.186 | 0.196 | 0.739 | 0.197 | 0.220 | 0.194 |
| | multiplet | 0.0 | 0.0 | 0.0 | 0.0 | 0.0 | 0.0 |
| | nc | 0.0 | 0.0 | 0.0 | 0.0 | 0.0 | 0.0 |
| **RNA-FM** | pk | 0.195 | 0.202 | 0.752 | 0.179 | 0.246 | 0.201 |
| | multiplet | 0.002 | 0.003 | 0.837 | 0.001 | 0.008 | 0.003 |
| | nc | 0.234 | 0.247 | 0.936 | 0.251 | 0.279 | 0.248 |
| **RiNALMo** | pk | 0.328 | 0.340 | 0.757 | 0.076 | 0.339 | 0.345 |
| | multiplet | 0.0 | 0.0 | 0.0 | 0.0 | 0.0 | 0.0 |
| | nc | - | - | - | - | - | - |
| **ProtRNA** | pk | 0.330 | 0.340 | 0.785 | 0.336 | 0.372 | 0.345 |
| | multiplet | 0.014 | 0.015 | 0.831 | 0.011 | 0.025 | 0.019 |
| | nc | 0.258 | 0.272 | 0.935 | 0.258 | 0.325 | 0.273 |

**Table S5. Secondary structure prediction comparison on inter-family benchmark dataset for pseudoknots (pk), multiplets, and noncanonical pairs (nc).** '-' indicates that the model either does not support predicting these interactions by design or has this feature disabled by default.

| Inter-Family Benchmark | | **F1** | **MCC** | **Weisfeiler Lehman** | **Recall** | **Precision** | **F1-shifted** |
|---|---|---|---|---|---|---|---|
| **SPOT-RNA** | pk | 0.185 | 0.245 | 0.684 | 0.155 | 0.261 | 0.190 |
| | multiplet | 0.099 | 0.104 | 0.801 | 0.090 | 0.137 | 0.120 |
| | nc | 0.239 | 0.274 | 0.929 | 0.183 | 0.464 | 0.261 |
| **MXFold2** | pk | - | - | - | - | - | - |
| | multiplet | 0.0 | 0.0 | 0.0 | 0.0 | 0.0 | 0.0 |
| | nc | - | - | - | - | - | - |
| **UFold** | pk | 0.037 | 0.042 | 0.673 | 0.027 | 0.073 | 0.042 |
| | multiplet | 0.0 | 0.0 | 0.0 | 0.0 | 0.0 | 0.0 |
| | nc | 0.0 | 0.0 | 0.0 | 0.0 | 0.0 | 0.0 |
| **RNA-FM** | pk | 0.112 | 0.115 | 0.650 | 0.099 | 0.148 | 0.120 |
| | multiplet | 0.012 | 0.014 | 0.828 | 0.008 | 0.030 | 0.020 |
| | nc | 0.129 | 0.149 | 0.852 | 0.095 | 0.263 | 0.180 |
| **RiNALMo** | pk | 0.230 | 0.234 | 0.594 | 0.23 | 0.263 | 0.253 |
| | multiplet | 0.0 | 0.0 | 0.0 | 0.0 | 0.0 | 0.0 |
| | nc | - | - | - | - | - | - |
| **ProtRNA** | pk | 0.100 | 0.110 | 0.692 | 0.076 | 0.179 | 0.105 |
| | multiplet | 0.031 | 0.033 | 0.814 | 0.029 | 0.046 | 0.052 |
| | nc | 0.160 | 0.192 | 0.857 | 0.120 | 0.356 | 0.201 |

**Table S6. Mean True Positive Rate (TPR) and False Positive Rate (FPR) for Protein-RNA interaction prediction of different language models and comparative methods across 17 RBPs in Hela cells.**

| | TPR ↑ | FPR ↓ | TPR-FPR ↑ |
|---|---|---|---|
| RealSS + Seq | 0.8031 | 0.1511 | 0.6520 |
| ProtRNA + Seq | 0.8171 | 0.1684 | 0.6486 |
| RiNALMo + Seq | 0.8086 | 0.1682 | 0.6404 |
| RNA-FM + Seq | 0.8126 | 0.1923 | 0.6204 |
| RNAErnie + Seq | 0.8072 | 0.1852 | 0.6220 |
| Seq | 0.8153 | 0.1800 | 0.6352 |

**Table S7. MRL prediction performance evaluation on 5'UTR sequences from Random7600 and Human 7600 dataset.**

| Model | **Random7600** | | **Human7600** | |
|---|---|---|---|---|
| | $R^2$ | MAE | $R^2$ | MAE |
| RNA-FM | 0.5498 | 0.6801 | 0.6074 | 0.5322 |
| RNAErnie | 0.5564 | 0.6823 | 0.6212 | 0.5274 |
| RiNALMo | 0.5717 | 0.6620 | 0.6307 | 0.5171 |
| ProtRNA | 0.5932 | 0.6402 | 0.6476 | 0.5030 |

**Table S8. Protein binary and subcellular localization prediction accuracy of ESM-2 and ProtRNA.**

| Model | **Binary Localization** | **Subcellular Localization** |
|---|---|---|
| ESM-2 | 0.916 | 0.802 |
| ProtRNA | 0.736 | 0.382 |
| LoRA | 0.726 | 0.416 |
